# A Deep Learning-Based Scoring Framework for Large-Scale Multi-Donor Cardiotoxicity Screening

**DOI:** 10.64898/2026.04.22.720197

**Authors:** Danny Vu, Andrew Kowalczewski, Sarah D. Burnett, Courtney Sakolish, Xiyuan Liu, Huaxiao Yang, Ivan Rusyn, Zhen Ma

**Affiliations:** Department of Biomedical & Chemical Engineering, Syracuse University, Syracuse, NY, USA; BioInspired Institute for Materials and Living Systems, Syracuse University, Syracuse, NY, USA; Department of Veterinary Physiology and Pharmacology, Texas A&M University, College Station, TX, USA; Department of Mechanical & Aerospace Engineering, Syracuse University, Syracuse, NY, USA; Department of Biomedical Engineering, University of North Texas, Denton, TX, USA

**Keywords:** Deep Learning, Cardiotoxicity, Human Induced Pluripotent Stem Cells, High-Throughput Screening

## Abstract

Cardiotoxicity remains a major cause of drug attrition and post-market withdrawal, yet the vast majority of environmental chemicals to which humans may be exposed remain uncharacterized for cardiotoxicity risk. Human induced pluripotent stem cell (hiPSC)-based testing has been proposed to address this gap. Here we present an unsupervised deep learning framework for multi-donor cardiotoxicity screening using high-throughput calcium transient recordings from hiPSC-derived cardiomyocytes (hiPSC-CMs). We used data from a library of 1,029 compounds that were tested in hiPSC-CM from five donors in concentration-response. An autoencoder trained exclusively on baseline signals quantified chemical-induced functional perturbations through reconstruction error, bypassing the need for labeled training data while capturing the full spectrum of calcium handling disruptions. Using this framework, we generated effect levels. Aggregation of donor-specific scores revealed substantial inter-individual variability in potential cardiotoxicity, underscoring the value of this approach for population-level risk prediction. We found that microbiocides, dyes, and pesticides to have potential concern, characterized by high toxicity scores and low inter-donor variability. This framework establishes a scalable, human-relevant, and genetically diverse platform for cardiotoxicity surveillance across both pharmacological and environmental chemical spaces, with direct implications for drug and chemical safety evaluation and prioritization for additional studies.

## INTRODUCTION

Cardiotoxicity remains a major cause of late-stage clinical failure and post-market drug withdrawal, accounting for approximately 14% of drugs removed due to adverse cardiac effects^1^. While pharmaceutical compounds have been the primary focus of safety evaluation, the challenge extends to environmental chemicals, which frequently undergo limited or insufficient cardiac safety evaluations but are known to exacerbate cardiovascular disease^2^. Over the past seven decades, industrial chemical production has increased more than twentyfold, introducing thousands of synthetic compounds into the environment^3^. Continuous human exposure occurs through air, water, food, and consumer products, yet the majority of these chemicals remain inadequately characterized for cardiac risk^4^. This disconnect underscores a critical unmet need for scalable, human-relevant experimental models capable of assessing cardiotoxic risk across both pharmacological and environmental chemical spaces, while accounting for the inter-individual variability.

Human induced pluripotent stem cell-derived cardiomyocytes (hiPSC-CMs) have emerged as a transformative platform for cardiotoxicity screening, offering scalability, human specificity, and compatibility with high-throughput functional assays^5-7^. Recent advances enable large-scale screening of drug responses from hiPSC-CMs in 96- and 384-well formats, with quantitative extraction of electrophysiological and contractile features such as peak amplitude, beat rate, and calcium decay^8-10^. These features have been increasingly integrated with machine learning models to enhance predictive modeling of drug-induced cardiotoxicity^11-15^. However, these approaches are predicated on deconvolution of the entire signal into individual features and thus rely on predefined metrics and may fail to capture higher-order temporal dynamics and subtle nonlinear patterns inherent in raw functional signals.

Deep learning approaches have begun to redefine signal analysis by directly operating on raw functional data, bypassing manual feature engineering and capturing subtle patterns that may be overlooked in feature extraction. Hybrid convolutional–recurrent architectures have been developed to detect arrhythmic irregularities in field potential recordings from hiPSC-CMs exposed to drugs^16^. A supervised convolutional neural network trained on optical voltage traces has achieved robust classification of drug-induced phenotypes including non-arrhythmic, arrhythmic, and asystolic states, while generating continuous, dose-dependent proarrhythmic risk metrics^17^. More recent studies have expanded these frameworks to integrate multimodal datasets, including high-content imaging and processed electrophysiological signals^18,19^. Despite these advances, current approaches remain constrained by limited chemical diversity, reliance on supervised classification paradigms requiring labeled training data, and insufficient representation of inter-individual variability across genetically distinct donors — thereby restricting their translational relevance to population-level risk prediction on environmental toxins.

Here, we present an unsupervised multi-donor deep learning framework and apply it to the data from a large-scale (1,000+ compounds tested in concentration-response) cardiotoxicity screening that used high-throughput calcium transient recordings from hiPSC-CMs derived from five human donors. First, an autoencoder trained exclusively on healthy baseline signals was used to quantify chemical-induced functional perturbations through reconstruction error, bypassing the need for labeled training data and enabling sensitivity to a broad spectrum of calcium handling disruptions. Next, the approach was applied to the data from a large library of 1,029 compounds that included pharmaceuticals with known cardiotoxicity risk classifications and environmental chemicals from the U.S. EPA ToxCast program representing 13 chemical classes^20^ (Figure 1). These analyses were performed on the data from each of five donors, and donor-specific reconstruction error scores were aggregated into a unified multi-donor voting metric to discover multi-donor dose–response relationships. Safety margins derived from this multi-donor scoring framework were benchmarked against existing deep learning frameworks and hERG-based approaches. Donor-specific vulnerability analyses revealed substantial inter-individual differences in drug effects, with microbiocides, dyes, and pesticides emerging as the chemical classes of greatest potential cardiotoxic concern. Together, this framework provides a scalable, human-relevant, and donor-aware strategy for systematic cardiotoxicity risk assessment for both drugs and environmental chemicals.

**Figure 1.**
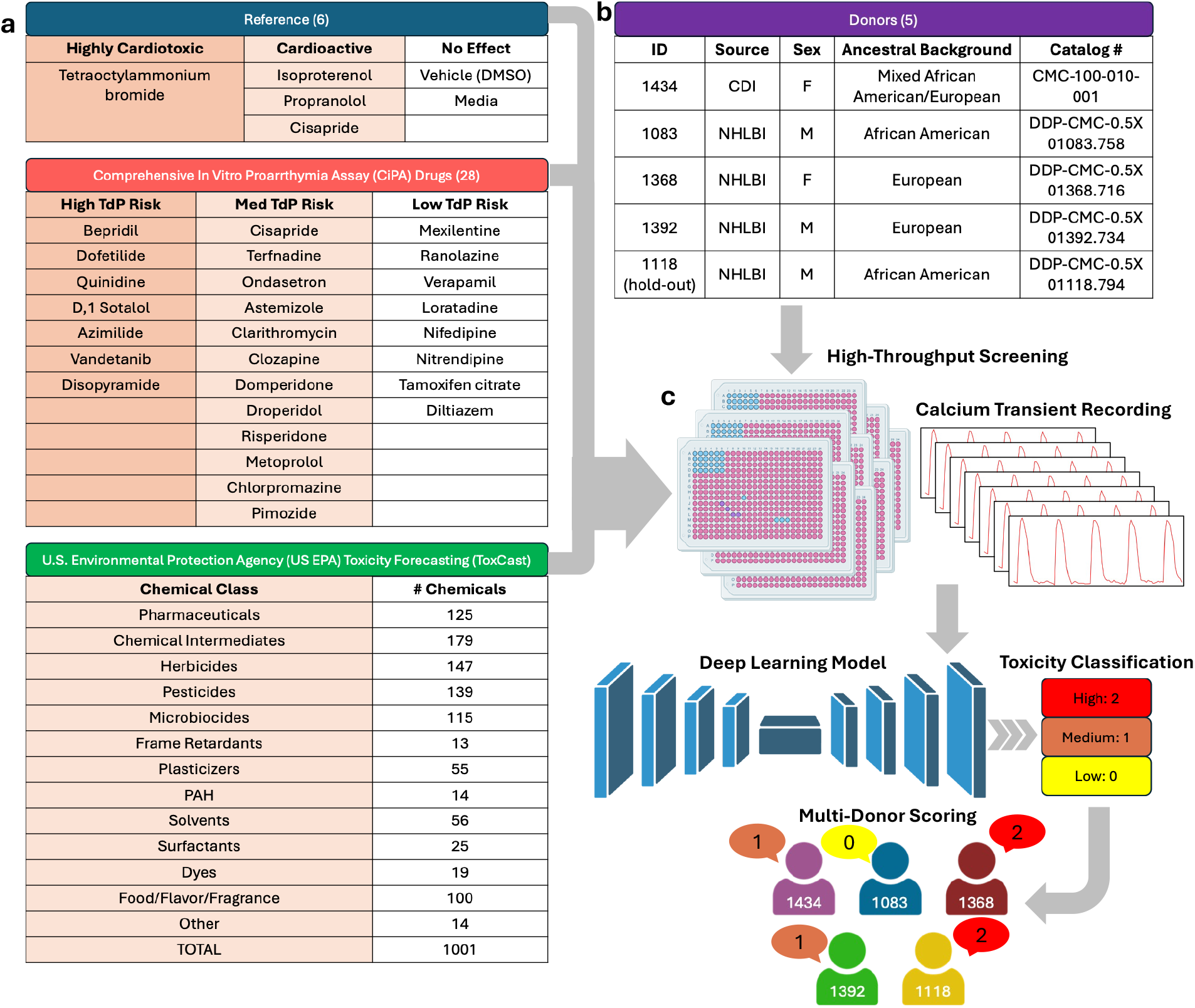
Overview of the multi-donor high-throughput cardiotoxicity screening framework. The chemical library comprised six reference conditions and 1,029 compounds (28 CiPA drugs and U.S. EPA ToxCast program representing 13 chemical classes). Compounds were screened in a 384-well format across five hiPSC-CM lines derived from genetically distinct donors. Calcium transient recordings were processed through an autoencoder-based deep learning model, which assigned each signal a reconstruction error-based risk score of 0 (low), 1 (medium), or 2 (high). Scores were aggregated across donors to generate a unified multi-donor cardiotoxicity metric.

## RESULTS

### Autoencoder Model Quantifies Chemically Induced Signal Perturbations

The data used for these studies was previously published^20^. The screening library comprised 1,029 compounds spanning three chemical categories (Figure 1a): six reference compounds with well-characterized cardioactive profiles, 28 Comprehensive in vitro Proarrhythmia Assay (CiPA) initiative drugs with defined torsades de pointes (TdP) risk classifications, and environmental chemicals from the U.S. EPA ToxCast program representing 13 chemical classes, including pesticides, plasticizers, flame retardants, surfactants, etc. Compounds were screened in a 384-well format with reference compounds included on each plate to benchmark assay performance and establish cardiotoxicity classification thresholds. Calcium transient recordings were acquired for 100 seconds at baseline and 90 minutes post-exposure across four concentrations (0.1, 1, 10, and 100 μM) from five hiPSC-CM lines derived from genetically distinct donors (Figure 1b), yielding a total dataset of 38,808 kinetic recordings. These recordings were processed through an autoencoder-based deep learning model, which assigned each signal a reconstruction error-based risk score of 0 (low toxicity), 1 (medium toxicity), or 2 (high toxicity). Scores were then aggregated across donors on a per-compound, per-concentration basis to generate a unified multi-donor cardiotoxicity metric (Figure 1c). To rigorously evaluate model generalizability, donor 1118 was withheld entirely from model training and reserved for independent validation, enabling a leave-one-donor-out assessment of inter-individual variability in cardiotoxic effects.

We developed an autoencoder-based deep learning framework comprising a symmetric encoder-decoder architecture with five fully connected layers in each branch (Figure 2a). The encoder progressively compressed 400-timepoint input signals through hidden layers into a 12-node latent space, while the decoder reconstructs the original signal. All hidden layers used hyperbolic tangent (tanh) activation functions to capture nonlinear signal dynamics, with dropout layers (rate = 0.2) applied after each dense layer to improve generalization. The model was trained exclusively on 19,404 baseline calcium transient recordings from four donors (1434, 1083, 1368, and 1392), enabling it to learn the intrinsic functional signatures of healthy hiPSC-CMs (Figure 2b). The autoencoder successfully reconstructed the baseline signals. Loss curves demonstrated rapid initial convergence over 500 epochs, with training loss (MSE ≈ 0.0048) and validation loss (MSE ≈ 0.0053) remaining closely coupled throughout training, indicating effective generalization without overfitting (Supplemental Figure 1a). To further assess model robustness, 5-fold cross-validation was performed on the baseline training set, yielding highly consistent MSE values across all folds, confirming stable reconstruction performance across held-out data partitions (Supplemental Figure 1b). The reconstructed signals closely matched with original baseline calcium transient signals (Supplemental Figure 1c)

**Figure 2.**
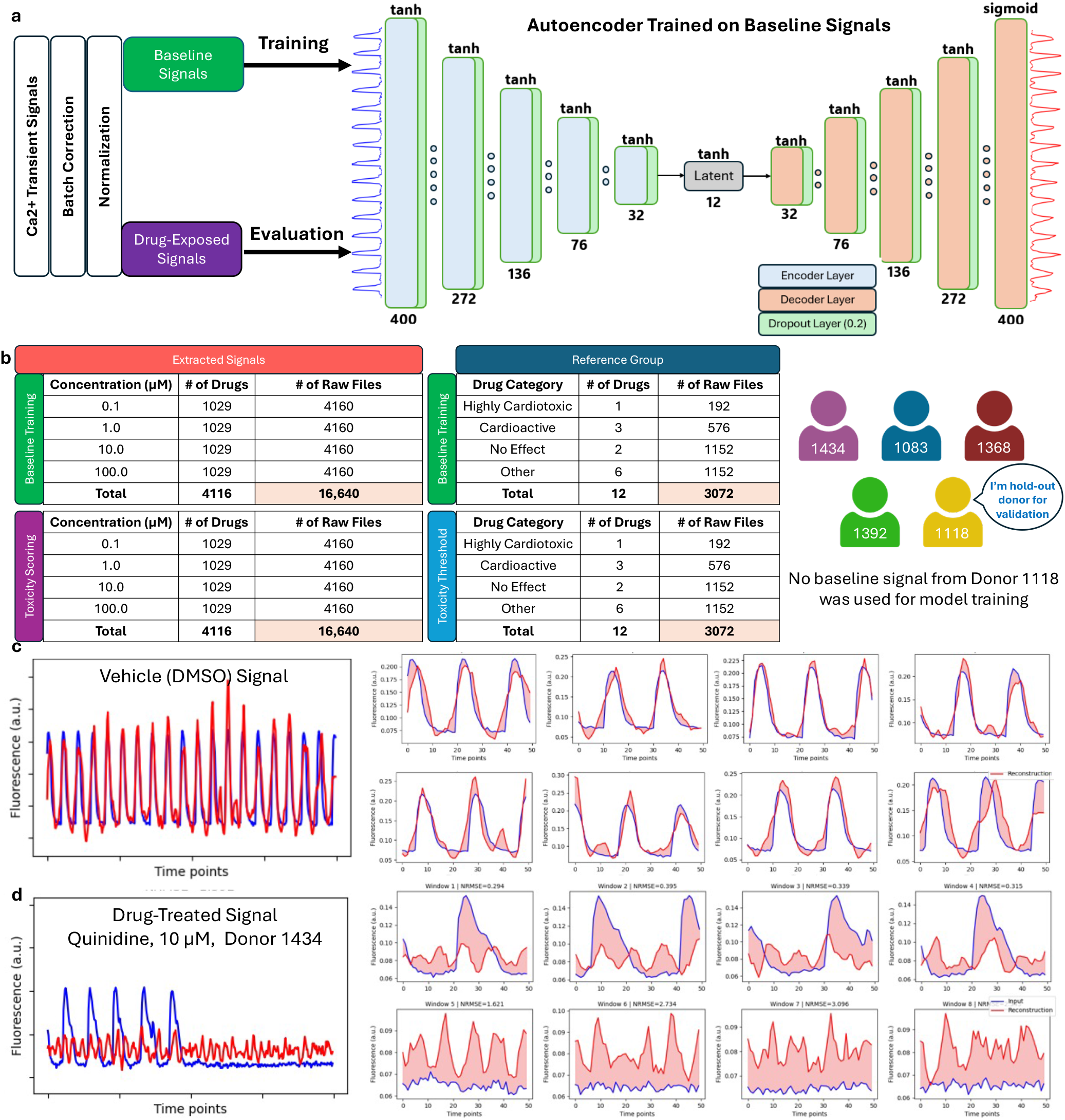
Autoencoder architecture for reconstruction error quantification. (a) Schematic of the symmetric encoder-decoder architecture comprising five fully connected layers in each branch. (b) Dataset design with baseline signals for model training (chemical treatment and reference groups), drug-exposed signals from reference groups for toxicity thresholding, and drug exposed signal from chemical treatment group for toxicity scoring. Donor 1118 designated as the held-out validation donor. (c) Representative reconstruction of a vehicle (DMSO)-treated calcium transient signal, showing close agreement between the original (blue) and reconstructed (red) traces across all eight windowed segments. (d) Representative reconstruction of a drug-treated calcium transient, demonstrating pronounced deviations between original and reconstructed traces, reflecting disrupted calcium handling dynamics.

Following training, the autoencoder was applied to calcium transients acquired after chemical exposure. Because the model was trained exclusively on baseline signals, it exhibited limited capacity to accurately reconstruct chemically perturbed signals, resulting in pronounced deviations between reconstructed outputs and original inputs. Reconstruction errors were quantified using a windowed normalized root mean square error (NRMSE), which normalizes error relative to the dynamic range of each signal. The autoencoder effectively reconstructed calcium transients from vehicle-treated controls, yielding low NRMSE values (Figure 2c). In contrast, chemically perturbed signals with pronounced chronotropic effects and altered peak-to-peak intervals produced substantially elevated NRMSE values, reflecting disrupted calcium handling dynamics (Figure 2d). While reconstruction error provided a continuous measure of deviation from baseline calcium dynamics, interpretation of these values required a structured framework to distinguish biologically meaningful drug effects.

### Multi-Donor Score Aggregation Enables Dose–Response Estimation

To stratify drug-exposed signals into cardiotoxicity risk categories, NRMSE-based thresholds were empirically derived from reference compounds with well-characterized cardioactive profiles (Figure 3a). Three reference groups were defined: a high-risk group comprising a cytotoxic compound tetraoctylammonium bromide (TAB, 50µM), high-TdP risk CiPA drugs at 100 µM concentration, a cardioactive group including Isoproterenol (10 µM), Propranolol (0.5 µM), and Cisapride (0.1 µM), and a no-effect group consisting of vehicle and media controls (Supplemental Figure 2). The high-medium threshold was set at the midpoint between the lowest NRMSE in the high-risk group and the highest NRMSE in the cardioactive group, yielding a value of 0.582. The medium-low threshold was defined as the midpoint between the mean NRMSE values of the cardioactive and no-effect groups, yielding a value of 0.291. Signals were then assigned scores of 2 (high toxicity), 1 (medium toxicity), or 0 (low toxicity) based on these thresholds, and scores were aggregated across four donors per compound per concentration, yielding a maximum possible score of 8 (Figure 3b). Applying this framework to all drug-exposed signals, the majority were classified as low toxicity (8,357) or medium toxicity (7,561), with a smaller subset categorized as high toxicity (3,486) (Figure 3c).

**Figure 3.**
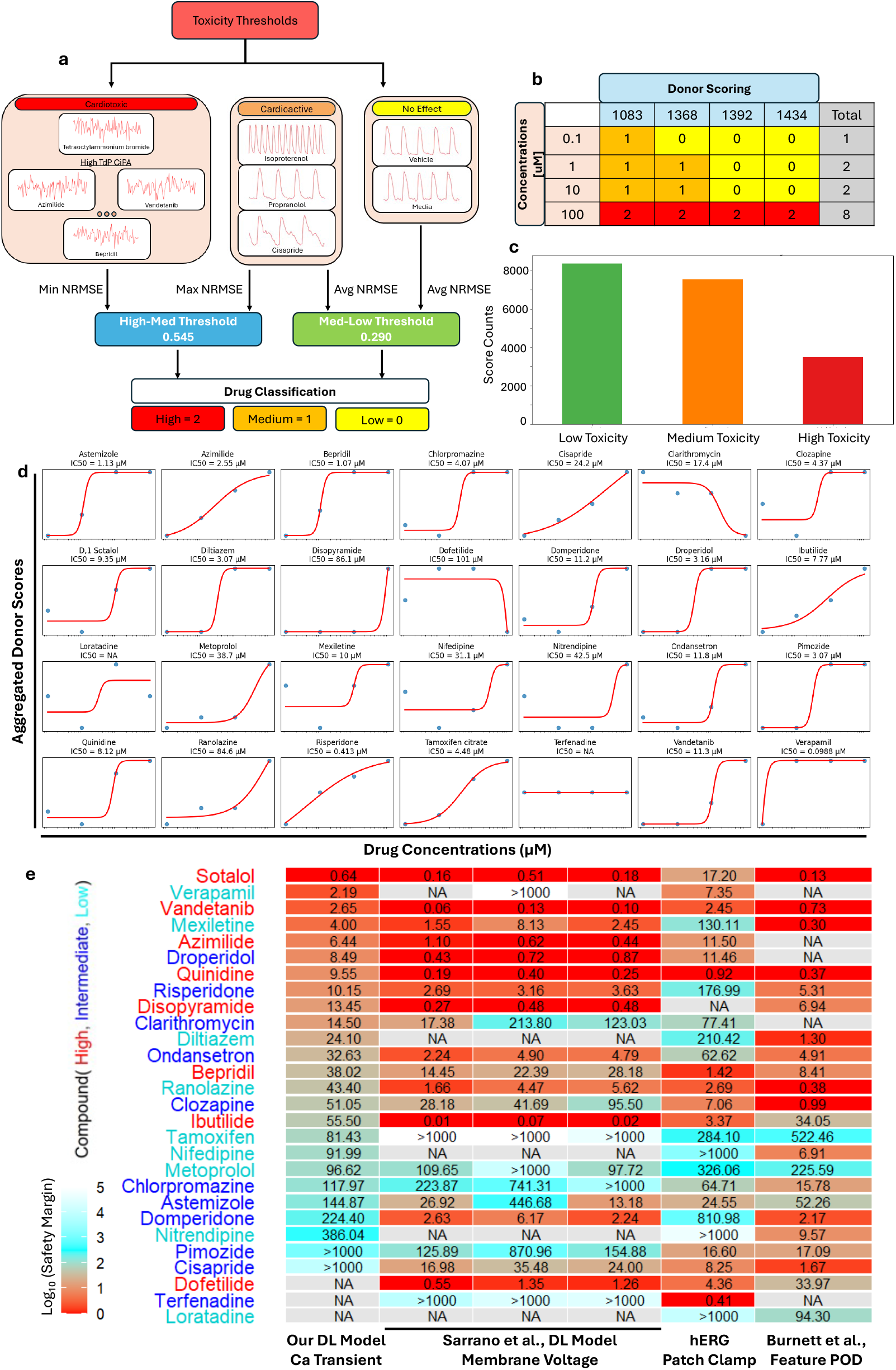
Multi-donor cardiotoxicity scoring for dose–response analysis on CiPA compounds. (a) Reference groups used to derive NRMSE-based cardiotoxicity thresholds for cardiotoxicity classification. (b) Representative donor scoring table illustrating concentration-dependent score aggregation across four donors. (c) Distribution of toxicity classifications across all drug-exposed signals. (d) Multi-donor dose–response curves for 24 of 28 CiPA drugs with estimable IC50 values. (e) Safety margin comparison between our deep-learning model, Serrano deep-learning model, hERG-based approaches, and Burnett feature-based POD model.

Aggregation of donor-specific scores across concentrations enabled construction of multi-donor dose–response relationships. Among the 1,029 compounds tested, 840 exhibited well-defined monotonic dose–response curves suitable for IC50 estimation using the Hill equation. The remaining 193 compounds displayed one of three atypical response patterns: minimal response across all concentrations, non-monotonic dose dependence, or pronounced donor-specific variability that precluded reliable curve fitting. Of the 28 CiPA drugs tested, we were able to obtain IC50 values for 25 of them (Figure 3d and Supplemental Table 1). Three compounds (Dofetilide, Terfenadine, and Loratadine) did not produce well-defined monotonic responses and were assigned IC50 = NA. Verapamil exhibited the lowest IC50 (0.1 μM), followed by Bepridil (1.07 μM), consistent with their potent disruption of cardiomyocyte electrophysiology. Medium-TdP risk compounds showed a broader range of IC50 values, from Droperidol (3.16 μM) to Metoprolol (38.7 μM), reflecting heterogeneous mechanisms of cardioactive perturbation. Low-risk compounds including Nifedipine (31 μM) and Nitrendipine (42.5 μM) generally exhibited higher IC50 values.

To assess the translational relevance of autoencoder-derived IC50 estimates, safety margins were compared against values reported by Serrano et al. deep-learning model^17^, Burnett et al. using feature-based point-of-departure (POD) method^20^, and traditional hERG-based measurements^21^ (Figure 3e). Compounds with smaller safety margins indicate a narrower therapeutic window, reflecting a higher likelihood of cardiotoxic effects at clinically relevant concentrations. High TdP risk compounds demonstrated the strongest overall concordance, with consistently low safety margins across all approaches. Safety margins derived from Burnett et al. were consistently low across many compounds, likely reflecting their framework’s use of conservative POD thresholds based on the earliest detectable phenotypic perturbation. Sotalol, Azimilide, Quinidine, and Vandetanib yielded small IC50 values in our autoencoder framework, concordant with their established high-risk classifications and generally low Serrano estimates. In comparison, Verapamil showed higher agreement between our model and hERG, while Serrano’s model reported the safety margin >1000 for one donor, likely reflecting differences in the functional endpoints captured — calcium transient reconstruction error being inherently sensitive to L-type calcium channel blockade, which is Verapamil’s primary mechanism of action.

Medium TdP risk compounds exhibited the greatest variability across frameworks, consistent with the mechanistic heterogeneity of this category. Ibutilide showed large discrepancy, with the autoencoder yielding the lowest safety margin (55.50), compared to low margin from the Serrano models (0.01 – 0.07) and hERG-based value (3.37). Low TdP risk compounds showed better concordance across frameworks, predominantly exhibiting large IC50 values (e.g. Nitrendipine and Metoprolol) or NA designations (e.g., Loratadine,) for compounds with minimal functional perturbation at tested concentrations. Overall, concordance between the autoencoder and orthogonal frameworks was strongest for high TdP risk compounds, where potent functional disruption produces consistent signals regardless of the assay modality employed. Differences in safety margin estimates across frameworks reflect their distinct mechanistic sensitivities — calcium transient-based reconstruction error captures perturbations in intracellular calcium dynamics, voltage-based endpoints reflect membrane potential changes, and hERG-based margins are limited to potassium channel blockade.

Next, we plotted safety margins across chemicals in our dataset with available exposure data, where pharmaceuticals were evaluated using population median *C*_*max*_ values and environmental chemicals were evaluated using the median steady-state plasma concentrations derived from ExpoCast exposure estimates and toxicokinetic modeling (Supplemental Figure 3). Pharmaceutical compounds exhibited the lowest safety margins of all groups, with median log10 values below 2. In contrast, environmental chemical classes exhibited systematically a wide range of safety margins, which is comparable to the original Burnett et al. analysis. Taken together, these findings demonstrate that while pharmaceutical compounds pose the most immediate cardiotoxic risk relative to exposure, several environmental chemical classes — particularly solvents, surfactants, and a subset of pesticides — warrant heightened surveillance due to their comparatively narrow safety margins.

### Donor Cardiotoxicity Vulnerability Across the ToxCast Library

Aggregated toxicity scores across ToxCast chemical classes revealed distinct risk profiles among environmental compound categories (Figure 4a). Microbiocides exhibited the highest proportion of high-toxicity responses (30.9%), followed by dyes (25.3%) and pesticides (23.8%), while solvents showed the lowest high-toxicity fraction (7.4%) and the highest proportion of low-toxicity responses (53.9%). Pharmaceuticals and flame retardants occupied intermediate positions, with high-toxicity proportions of 23.4% and 17.7%, respectively. These patterns were largely consistent with known biological activity profiles that microbiocides and pesticides are designed to disrupt cellular function.

**Figure 4.**
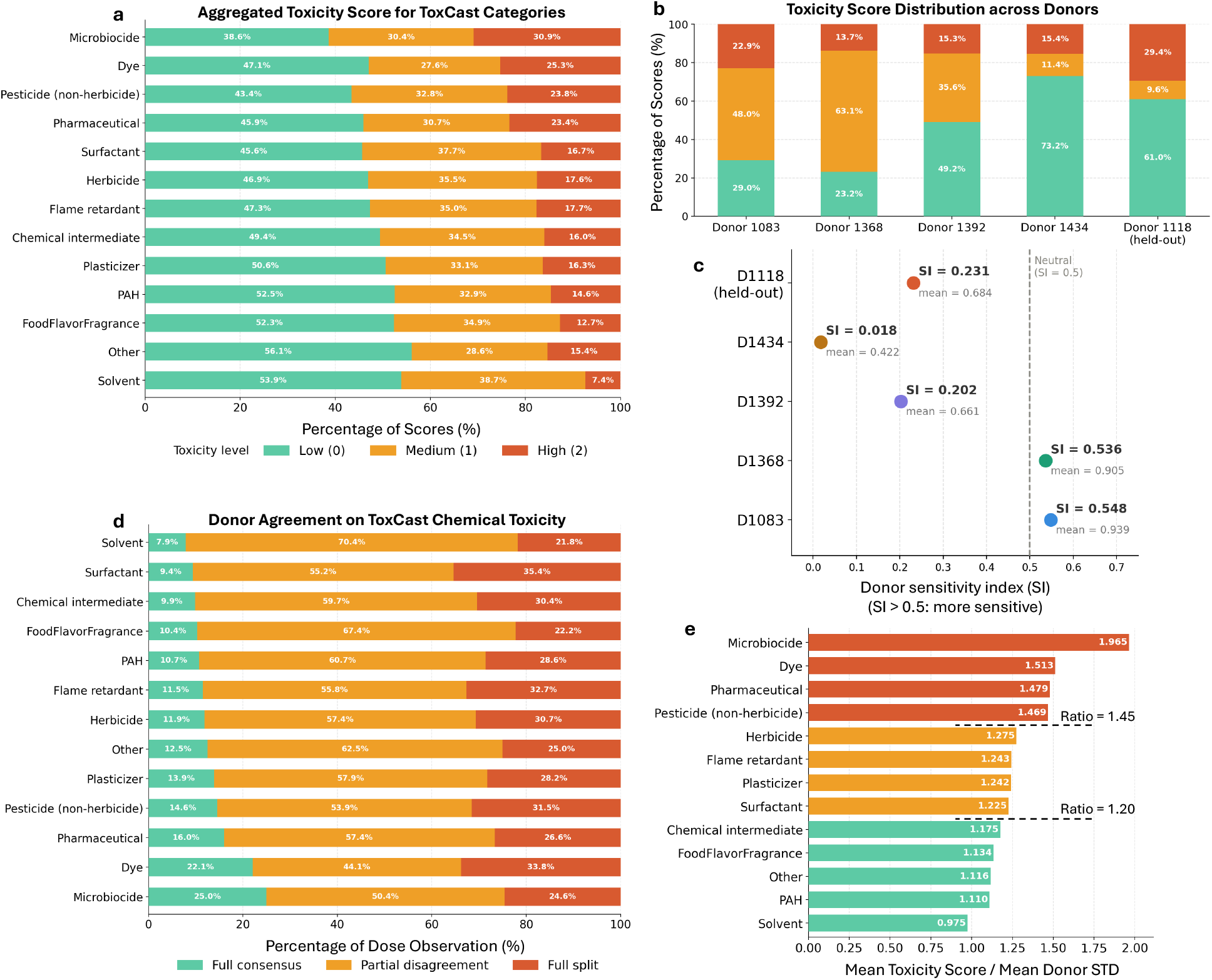
Donor-specific vulnerability and cardiotoxicity profiling for ToxCast chemical classes. (a) Aggregated toxicity score distributions across ToxCast chemical classes, showing microbiocides and dyes as the highest toxicity categories. (b) Donor differences in signal classification into low, medium, and high toxicity, including the held-out donor 1118. (c) Scatter plot of sensitivity index (SI) versus mean NRMSE for each donor. The neutral threshold (SI = 0.5) is shown as a dashed reference line. (d) Donor agreement analysis across chemical classes, showing the percentage of dose observations classified as full consensus, partial disagreement, or full split. Solvents and surfactants showed the least donor consensus. (e) Risk ratio (mean toxicity score / mean donor STD) for each ToxCast chemical class, ordered by descending ratio. Dashed lines at 1.45 and 1.20 delineate high-concern, moderate-concern, and low-concern tiers. Microbiocides exhibited the highest ratio (1.965), indicating consistently elevated cardiotoxic effects with low inter-donor variability.

Toxicity score distributions varied substantially across donors, revealing clear differences in cardiomyocyte sensitivity to chemical exposure (Figure 4b and Supplemental Figure 4). Donor 1083 and 1368 exhibited the highest susceptibility, with combined medium and high toxicity fractions of 70.9% and 76.8% and vulnerability indices of SI = 0.548 and SI = 0.536, respectively (Figure 4c). In contrast, donor 1434 displayed the greatest resistance to chemical effects, with 73.2% of signals classified as low toxicity and the lowest vulnerability index (SI = 0.018, mean NRMSE = 0.422). The held-out donor 1118 exhibited the largest high-toxicity fraction among all donors (29.4%), with a sensitivity index of 0.231 and mean NRMSE of 0.684, positioning it between donors 1392 and 1368 in overall vulnerability. The model successfully captured the elevated cardiotoxic response profile of this unseen donor, whose vulnerability fell within the range spanned by the training donors, demonstrates the robustness and generalizability of the reconstruction error framework across genetically diverse hiPSC-CM lines. Different donors showed variable responses to the ToxCast chemicals from different classes (Supplemental Figure 5). Analysis of donor agreement across chemical classes demonstrated that consensus responses, where all donors agreed on toxicity classification, were most prevalent for solvents (70.4% full consensus) and surfactants (55.2%). Microbiocides showed the lowest full consensus (25.0%) and the highest rate of full donor disagreement (24.6%) (Figure 4d), suggesting that individual genetic background substantially influences cardiomyocyte vulnerability to this chemical class.

To identify chemical classes posing the greatest cardiotoxic concern, the ratio of mean toxicity score to mean donor standard deviation (STD) was calculated for each ToxCast category (Figure 4e). A high ratio indicates that a chemical class produces consistently elevated cardiotoxic responses across all donors, representing the highest concern for broad human health risk from environmental exposure. Microbiocides exhibited the highest ratio (1.965), followed by dyes (1.513), pharmaceuticals (1.479), and pesticides (1.469), all exceeding the high-concern threshold. In contrast, PAHs (1.110), food/flavor/fragrance compounds (1.134), and solvents (0.975) exhibited the lowest ratios, with solvents falling below 1.0, indicating that their modest cardiotoxic signals were accompanied by relatively high inter-donor variability. To further resolve compound-level risk profiles within the ToxCast library, individual compounds were projected onto a mean toxicity score versus mean donor STD quadrant map (Supplemental Figure 6). Compounds in the upper-right quadrant represented high-risk (high toxicity scores) and high-variability (high donor STD) compounds with heterogeneous susceptibility, which is particularly relevant for environmental risk assessment, where exposure occurs across genetically diverse human populations Notable compounds in this category included Chlorpyrifos-methyl, MEHP, Simvastatin, 1,2-Dimethyl-4-nitrobenzene, Fenthion, and Myclobutanil, spanning pesticide, plasticizer, and pharmaceutical chemical classes. This might urge targeted follow-up studies using expanded donor panels and genetic sequencing to characterize the biological basis of their donor-dependent cardiotoxic responses.

## DISCUSSION

### Unsupervised Reconstruction Error as a Functional Cardiotoxicity Metric

In this study, we developed an unsupervised multi-donor deep learning framework for large-scale cardiotoxicity screening, which contrasts with previously reported supervised deep learning approaches that require predefined cardiotoxicity labels for model training^16,17^. By learning the intrinsic functional signatures of healthy cardiomyocytes from baseline recordings alone, the autoencoder generates a continuous, label-free measure of deviation from normal calcium dynamics. This approach bypasses the need for labeled training data while remaining sensitive to a broad spectrum of calcium handling disruptions. The biological validity of reconstruction error as a cardiotoxicity metric is supported by its concordance with other functional measurements. Autoencoder-derived IC50 values for CiPA compounds showed consistent trends with reported Ca_V1.2_ patch-clamp potencies for overlapping compounds including Bepridil, Quinidine, and Verapamil, suggesting that the latent features learned by the autoencoder capture perturbations in calcium-dependent electrophysiology^22,23^ (Supplemental Table 2). Furthermore, safety margins derived from multi-donor dose–response curves demonstrated risk rankings concordant with both the Serrano deep learning framework and hERG-based approaches for high-TdP risk compounds. These findings position reconstruction error as a functionally comprehensive cardiotoxicity metric that complements rather than replaces existing approaches.

### Multi-Donor Aggregation Enables Cardiotoxicity Risk Assessment

Single-donor screening strategies risk conflating donor-specific sensitivity with compound-level cardiotoxicity. By aggregating reconstruction error-based scores across four genetically distinct donors, our scoring framework produces multi-donor dose–response relationships with IC50 estimates (Supplemental Table 3). The substantial inter-individual variability observed across donors underscores the importance of this approach. Donor 1083 and 1368 exhibited the highest vulnerability indices, with combined medium and high toxicity fractions exceeding 70%, while donor 1434 displayed marked resistance, with 73.2% of signals classified as low toxicity. These differences likely reflect underlying genetic variation in ion channel expression, calcium handling capacity, and metabolic processing of xenobiotics — factors known to contribute to inter-individual differences in drug sensitivity in clinical populations^24,25^. The successful application of the trained framework to the held-out donor 1118, whose vulnerability index fell within the range spanned by the training donors, demonstrates that the autoencoder learned generalizable representations of cardiotoxic perturbation rather than donor-specific signal characteristics.

### Differential Vulnerability Across Chemical Classes Informs Environmental Risk Prioritization

Beyond pharmaceutical cardiotoxicity, our study systematically profiled over a large-scale library of environmental chemicals from the U.S. EPA ToxCast program — a chemical space that remains largely uncharacterized for cardiac risk despite widespread human exposure (Supplemental Table 4). The chemical class-level risk ratio analysis identified microbiocides, dyes, and pesticides (1.469) as the highest-concern categories, characterized by both elevated mean toxicity scores and low inter-donor variability, suggesting that cardiotoxic risk from these chemical classes is not restricted to genetically susceptible subpopulations but may represent a broadly applicable hazard. Conversely, the identification of high-toxicity, high-variability compounds highlights a distinct category of environmental chemicals whose cardiac risk may be concentrated in specific subpopulations. The organophosphate pesticides Chlorpyrifos-methyl and Fenthion, and the plasticizer MEHP are among the most prevalent environmental contaminants in human biomonitoring studies^26,27^. Their high inter-donor variability suggests that standard population-averaged risk assessments would underestimate risk for sensitive subpopulations, thus complicating regulatory decision-making.

### Limitations and Future Directions

Our autoencoder was trained on calcium transient recordings from four donors, which represents a limited sampling of the genetic diversity present in human populations. Expansion to larger donor panels, particularly those enriched for individuals with known cardiac risk factors such as long QT syndrome variants or drug metabolism polymorphisms, would further improve the population-level representativeness of the framework. Additionally, while calcium transient recordings provide a sensitive readout of intracellular calcium dynamics, they capture a subset of the electrophysiological endpoints relevant to cardiotoxicity assessment. Integration with complementary modalities, such as multielectrode array field potential recordings or contractility measurements, would enable more comprehensive characterization of drug-induced cardiac phenotypes. Finally, the threshold-based classification system, while empirically derived from well-characterized reference compounds, introduces discrete boundaries into what is inherently a continuous biological spectrum — future work could explore probabilistic scoring approaches that more fully leverage the continuous nature of reconstruction error distributions.

## METHODS

### Experimental details

Full experimental details for the wet-lab experiments are detailed in a previously published study^20^. Briefly, human induced pluripotent stem cell–derived cardiomyocytes (hiPSC-CMs) from five healthy donors were used. Donors were selected to represent individuals with no known history of cardiovascular disease, enabling assessment of inter-individual variability in drug response. For instance, relative to the standard donor (1434), donor 1083 exhibited a higher baseline beat rate, while donor 1392 showed a lower beat rate. Cells from each donor were cultured in a 384-well plate format under standardized conditions. Cardiomyocytes were maintained in culture for approximately 14 days prior to experimentation, during which they formed confluent monolayers and exhibited spontaneous, synchronous beating.

Test compounds were administered to hiPSC-derived cardiomyocytes using a high-throughput 384-well plate format. Cells were exposed to each compound at four concentrations (0.1, 1, 10, and 100 µM) in a final vehicle concentration of 0.5% dimethyl sulfoxide (DMSO). Compounds were distributed across multiple plates with randomized layouts, and each plate contained an identical set of controls, including vehicle-only and media-only wells. Pharmacological references were incorporated to capture key cardiomyocyte responses. Positive references included Isoproterenol and Propranolol to represent increases and decreases in beat rate, respectively, as well as Cisapride to capture delayed repolarization. Tetraoctylammonium bromide (TAB) was included as a cytotoxic control (Supplemental Figure 7). These controls were used to define baseline and reference responses for downstream cardiotoxicity classification.

Calcium transient recordings were acquired at baseline and following drug treatment using the FLIPR Tetra Cellular Screening System. Calcium signals were recorded at a sampling rate of 8 frames per second for 100 seconds, yielding approximately 800 time points per recording. Following baseline acquisition, cells were exposed to test compounds and incubated for 90 minutes before post-treatment recordings were obtained.

### Data Preprocessing

All calcium transient recordings were truncated to the middle 400 time points to standardize input length and reduce edge artifacts, and subsequently scaled to the range [0, 1] using min-max normalization to ensure consistent signal amplitudes across recordings and stable model training. To ensure the model learned physiologically representative calcium transient patterns, recordings with no detectable signal were removed prior to training. A percentile-based amplitude threshold derived from the baseline signal distribution was applied, with the 10th percentile used as the lower cutoff to exclude low-amplitude, noisy recordings. To reduce donor-specific variability in baseline calcium transient signals, batch correction was performed using the ComBat algorithm implemented in the Scanpy package. ComBat applies an empirical Bayes framework to adjust for systematic differences in signal distributions across donors by normalizing both the mean and variance and was applied to baseline recordings.

### Autoencoder Deep-Learning Model

The autoencoder architecture was created and optimized using the KerasTuner package in Python. The final model consisted of a symmetric encoder-decoder structure with five fully connected layers in each branch. The encoder progressively compressed 400-dimensional input signals through hidden layers of 272, 136, 76, and 32 nodes into a 12-node latent representation, capturing the most salient features of healthy calcium transient morphology. The decoder mirrored this structure, expanding the latent representation back through layers of 32, 76, 136, and 272 nodes to reconstruct the original 400-dimensional signal. All hidden layers employed hyperbolic tangent (tanh) activation functions to capture nonlinear signal dynamics, and dropout layers with a rate of 0.2 were applied after each dense layer in both the encoder and decoder to reduce overfitting and improve generalization across donors. The output layer used a sigmoid activation function to constrain reconstructed values to the [0, 1] range, consistent with the min-max scaled input data.

The autoencoder was trained using mean squared error (MSE) loss to encourage accurate reconstruction of baseline calcium transients. Training was performed for up to 500 epochs with a batch size of 64 and shuffled input data. To prevent overfitting and ensure stable convergence, two adaptive training strategies were employed: early stopping restored the best model weights if validation loss failed to improve for 20 consecutive epochs, and a learning rate scheduler reduced the learning rate by a factor of 0.5 after 10 epochs without improvement. Model performance was evaluated using training and validation loss curves to confirm stable convergence without overfitting. To further assess generalizability, 5-fold cross-validation was performed on the baseline training set, with MSE calculated separately for each fold to confirm consistent reconstruction performance across held-out data partitions.

### Reconstruction Error

Reconstruction error was quantified using windowed normalized root mean square error (NRMSE), which scales reconstruction error relative to the dynamic range of each individual signal:

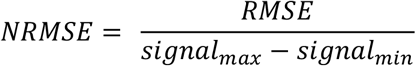

Each recording was divided into eight consecutive non-overlapping windows, and the NRMSE was calculated independently for each window. The final reconstruction error for each signal was computed as the mean NRMSE across all eight windows, providing a temporally resolved measure of deviation from learned baseline calcium dynamics.

### Cardiotoxicity Threshold

To translate reconstruction error measurements into biologically interpretable risk categories, NRMSE-based thresholds were empirically derived from reference compounds with well-characterized effects on hiPSC-CM calcium dynamics. Three risk categories were defined: a low-risk category consisting of vehicle-treated or media-control cells; a medium-risk category comprising compounds that significantly alter calcium transient features, specifically Isoproterenol (10 µM), Propranolol (0.5 µM), and Cisapride (0.1 µM); and a high-risk category encompassing compounds of tetraoctylammonium bromide (TAB, 50 µM) and CiPA high-risk drugs at the highest tested concentration (100 μM). These compounds were selected to establish benchmark cardiomyocyte responses for comparison with chemical-induced effects. Isoproterenol and Propranolol represent positive and negative chronotropy, respectively, while Cisapride captures changes in repolarization through the decay-rise ratio. TAB serves as a control for cytotoxicity, representing severe disruption of cardiomyocyte function.

Prior to threshold calculation, outlier signals were removed using the interquartile range (IQR) method to ensure robust estimation of representative reconstruction errors for each reference compound. Signals with NRMSE values exceeding 1.5 times the IQR above the third quartile or below the first quartile were excluded. The high-medium threshold was then calculated as the midpoint between the minimum NRMSE of the high-risk category and the maximum NRMSE of the medium-risk category, yielding a value of 0.545. The medium-low threshold was calculated as the midpoint between the mean NRMSE values of the medium-risk and low-risk categories, yielding a value of 0.290. Based on these thresholds, each drug-exposed signal was assigned a toxicity score of 2 (high risk), 1 (medium risk), or 0 (low risk), and scores were aggregated across donors per compound per concentration to yield a maximum possible score of 8.

### IC50 and Safety Margin Estimation

IC50 values were estimated by fitting the multi-donor aggregated dose–response data to the Hill equation using the SciPy optimization package in Python. Safety margins were calculated for CiPA compounds with well-defined dose–response curves as the ratio of the autoencoder-derived IC50 to the clinically relevant maximum plasma concentration (*C*_*max*_):

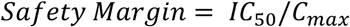

Compounds with smaller safety margins indicate a narrower therapeutic window and higher likelihood of cardiotoxic effects at clinically relevant concentrations. *C*_*max*_ values were obtained from published clinical pharmacokinetic data^28^.

### Donor Vulnerability Scoring

To quantify inter-individual differences in cardiomyocyte sensitivity to chemical exposure, a sensitivity index (SI) was calculated for each donor as a weighted ratio of toxicity classifications across all tested compounds:

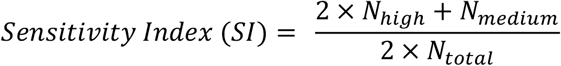

where *N*_*high*_, *N*_*medium*_, and *N*_*total*_ represent the number of high-toxicity, medium-toxicity, and total scored signals for each donor, respectively. High-toxicity responses were weighted twice as heavily as medium-toxicity responses to emphasize severe functional impairment. SI values range from 0 (complete resistance) to 1 (maximum vulnerability). Donor-specific toxicity score distributions were visualized as stacked bar charts showing the percentage of signals classified as low (0), medium (1), and high (2) toxicity for each donor. The relationship between SI and mean NRMSE across donors was visualized as a scatter plot, with each donor represented as a labeled point. The neutral SI threshold (SI = 0.5) was indicated as a reference line to distinguish relatively sensitive from relatively resistant donors.

### Chemical Class-Level Toxicity and Donor Agreement Analysis

To assess cardiotoxic responses across ToxCast chemical classes, toxicity scores were aggregated for all compounds within each chemical category. For each class, the percentage of signals classified as low, medium, and high toxicity was calculated across all donors and concentrations and visualized as a horizontal stacked bar chart ordered by descending high-toxicity fraction. Chemical class annotations were obtained from the U.S. EPA ToxCast database metadata.

Donor agreement was assessed at the per-compound, per-concentration level. For each condition, the distribution of donor toxicity scores was examined and classified into one of three agreement categories: full consensus, defined as all four donors assigned the same toxicity score; partial disagreement, defined as donors split across two adjacent toxicity categories; and full split, defined as donors distributed across all three toxicity categories or non-adjacent categories. The percentage of dose observations falling into each agreement category was calculated for each chemical class and visualized as a horizontal stacked bar chart ordered by descending full-consensus fraction.

To provide a summary metric for comparative cardiotoxic risk assessment across chemical classes, the ratio of the mean toxicity score to the mean donor standard deviation was calculated:

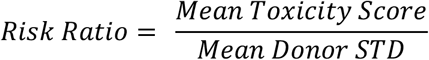

Risk ratios were visualized as a horizontal bar chart with chemical classes ordered by descending ratio value. Two reference thresholds were assigned at ratio values of 1.45 and 1.20 to delineate high-concern (ratio > 1.45), moderate-concern (1.20 < ratio ≤ 1.45), and low-concern (ratio ≤ 1.20) chemical categories.

### Compound-Level Toxicity and Variability Quadrant Analysis

The mean toxicity score, defined as the average of all donor toxicity scores, and the mean donor standard deviation (STD), defined as the average standard deviation of toxicity scores across, were calculated for each compound across all donors and concentrations. Compounds were plotted in a two-dimensional scatter plot with mean toxicity score on the x-axis and mean donor STD on the y-axis. The 75th percentile thresholds for both metrics (mean toxicity score = 0.90; mean donor STD = 0.63) were calculated across all compounds and overlaid as dashed reference lines to define four quadrants based on toxicity and variability.

## Supplemental Information

Supplemental information can be found online.

## Acknowledgement

This work was supported by the NIH [R01HD101130, R21HD11458, R15HD108720], the NSF [2130192 and 1943798].

## Author Contributions

D.V. and Z.M. conceived the study. D.V. and A.K. established the autoencoder deep learning model. D.V. established the multi-donor scoring framework. C. S. provided the dataset with detailed information. X.L. provided insights on machine learning-based analysis. H.Y. provided insights on AI models for cardiotoxicity evaluation. I.R. supervised the high-throughput screening of chemical cardiotoxicity. Z.M. supervised the AI model implementation and project development. D.V. and Z.M. wrote the manuscript with discussion and improvements from all authors. Z.M. funded the study.

## Resource Availability

The code for deep learning model for reconstruction error quantification of cardiomyocyte calcium transient is available at https://github.com/Danny-Vu/Syracuse-TAMU

Further information and requests for complete dataset of high-throughput chemical cardiotoxicity screening should be directed to and will be fulfilled by the contact, Zhen Ma.

## References

1 Onakpoya, I. J., Heneghan, C. J. & Aronson, J. K. Post-marketing withdrawal of 462 medicinal products because of adverse drug reactions: a systematic review of the world literature. BMC Med 14, 10 (2016). PMC4740994

2 Blaustein, J. R., Quisel, M. J., Hamburg, N. M. & Wittkopp, S. Environmental Impacts on Cardiovascular Health and Biology: An Overview. Circ Res 134, 1048–1060 (2024). PMC11058466

3 Woodruff, T. J., Burke, T. A. & Zeise, L. The need for better public health decisions on chemicals released into our environment. Health Aff (Millwood) 30, 957–967 (2011). PMC6709852

4 Maffini, M. V. & Vandenberg, L. N. Science evolves but outdated testing and static risk management in the US delay protection to human health. Front Toxicol 6, 1444024 (2024). PMC11347445

5 Magdy, T., Schuldt, A. J. T., Wu, J. C., Bernstein, D. & Burridge, P. W. Human Induced Pluripotent Stem Cell (hiPSC)-Derived Cells to Assess Drug Cardiotoxicity: Opportunities and Problems. Annu Rev Pharmacol Toxicol 58, 83–103 (2018). PMC7386286

6 Nair, P., Prado, M., Perea-Gil, I. & Karakikes, I. Concise Review: Precision Matchmaking: Induced Pluripotent Stem Cells Meet Cardio-Oncology. Stem Cells Transl Med 8, 758–767 (2019). PMC6646696

7 Burnett, S. D., Blanchette, A. D., Chiu, W. A. & Rusyn, I. Human induced pluripotent stem cell (iPSC)-derived cardiomyocytes as an in vitro model in toxicology: strengths and weaknesses for hazard identification and risk characterization. Expert Opin Drug Metab Toxicol 17, 887–902 (2021). PMC8363519

8 Arkin, M. R. et al. in Assay Guidance Manual (eds S. Markossian et al.) (2004).

9 Watanabe, H., Honda, Y., Deguchi, J., Yamada, T. & Bando, K. Usefulness of cardiotoxicity assessment using calcium transient in human induced pluripotent stem cell-derived cardiomyocytes. J Toxicol Sci 42, 519–527 (2017).

10 Sirenko, O. et al. Assessment of beating parameters in human induced pluripotent stem cells enables quantitative in vitro screening for cardiotoxicity. Toxicol Appl Pharmacol 273, 500–507 (2013). PMC3900303

11 Yang, H. et al. Deriving waveform parameters from calcium transients in human iPSC-derived cardiomyocytes to predict cardiac activity with machine learning. Stem Cell Reports 17, 556–568 (2022). PMC9039838

12 Yang, H. et al. Prediction of inotropic effect based on calcium transients in human iPSC-derived cardiomyocytes and machine learning. Toxicol Appl Pharmacol 459, 116342 (2023).

13 Pang, J. K. S. et al. Characterizing arrhythmia using machine learning analysis of Ca(2+) cycling in human cardiomyocytes. Stem Cell Reports 17, 1810–1823 (2022). PMC9391413

14 Lee, E. K. et al. Machine Learning of Human Pluripotent Stem Cell-Derived Engineered Cardiac Tissue Contractility for Automated Drug Classification. Stem Cell Reports 9, 1560–1572 (2017). PMC5829317

15 Kowalczewski, A. et al. Integrating nonlinear analysis and machine learning for human induced pluripotent stem cell-based drug cardiotoxicity testing. J Tissue Eng Regen Med 16, 732–743 (2022). PMC9719611

16 Golgooni, Z. et al. Deep Learning-Based Proarrhythmia Analysis Using Field Potentials Recorded From Human Pluripotent Stem Cells Derived Cardiomyocytes. IEEE Journal of Translational Engineering in Health and Medicine 7, 1–9 (2019).

17 Serrano, R. et al. A deep learning platform to assess drug proarrhythmia risk. Cell Stem Cell 30, 86–95 e84 (2023). PMC9924077

18 Zhu, Z. et al. Two-Dimensional Deep Learning Frameworks for Drug-Induced Cardiotoxicity Detection. ACS Sens 9, 3316–3326 (2024).

19 Grafton, F. et al. Deep learning detects cardiotoxicity in a high-content screen with induced pluripotent stem cell-derived cardiomyocytes. Elife 10 (2021). PMC8367386

20 Burnett, S. D., Blanchette, A. D., Chiu, W. A. & Rusyn, I. Cardiotoxicity Hazard and Risk Characterization of ToxCast Chemicals Using Human Induced Pluripotent Stem Cell-Derived Cardiomyocytes from Multiple Donors. Chem Res Toxicol 34, 2110–2124 (2021). PMC8762671

21 Davies, M. R. et al. Use of Patient Health Records to Quantify Drug-Related Pro-arrhythmic Risk. Cell Rep Med 1, 100076 (2020). PMC7659582

22 Crumb, W. J., Jr., Vicente, J., Johannesen, L. & Strauss, D. G. An evaluation of 30 clinical drugs against the comprehensive in vitro proarrhythmia assay (CiPA) proposed ion channel panel. J Pharmacol Toxicol Methods 81, 251–262 (2016).

23 Kramer, J. et al. MICE models: superior to the HERG model in predicting Torsade de Pointes. Sci Rep 3, 2100 (2013). PMC3696896

24 Pang, L. et al. Predicting oncology drug-induced cardiotoxicity with donor-specific iPSC-CMs-a proof-of-concept study with doxorubicin. Toxicol Sci 200, 79–94 (2024). PMC11199917

25 Grimm, F. A. et al. A human population-based organotypic in vitro model for cardiotoxicity screening. ALTEX 35, 441–452 (2018). PMC6231908

26 Fagundes, T. R., Coradi, C., Vacario, B. G. L., de Morais Valentim, J. M. B. & Panis, C. Global Evidence on Monitoring Human Pesticide Exposure. J Xenobiot 15 (2025). PMC12641724

27 Barr, D. B. & Angerer, J. Potential uses of biomonitoring data: a case study using the organophosphorus pesticides chlorpyrifos and malathion. Environ Health Perspect 114, 1763–1769 (2006). PMC1665422

28 Vicente, J. et al. Mechanistic Model-Informed Proarrhythmic Risk Assessment of Drugs: Review of the “CiPA” Initiative and Design of a Prospective Clinical Validation Study. Clin Pharmacol Ther 103, 54–66 (2018). PMC5765372

