## Supplemental Figures for "A Deep Learning-Based Scoring Framework for Large-Scale Multi-Donor Cardiotoxicity Screening"

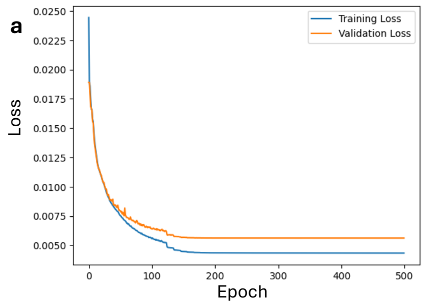


b

Mean Reconstruction Error across 5 Stratified Folds

| | Train | Test |
|---|---|---|
| **Fold 1** | 0.005469 | 0.005497 |
| **Fold 2** | 0.005479 | 0.005460 |
| **Fold 3** | 0.005482 | 0.005446 |
| **Fold 4** | 0.005474 | 0.005477 |
| **Fold 5** | 0.005470 | 0.005494 |

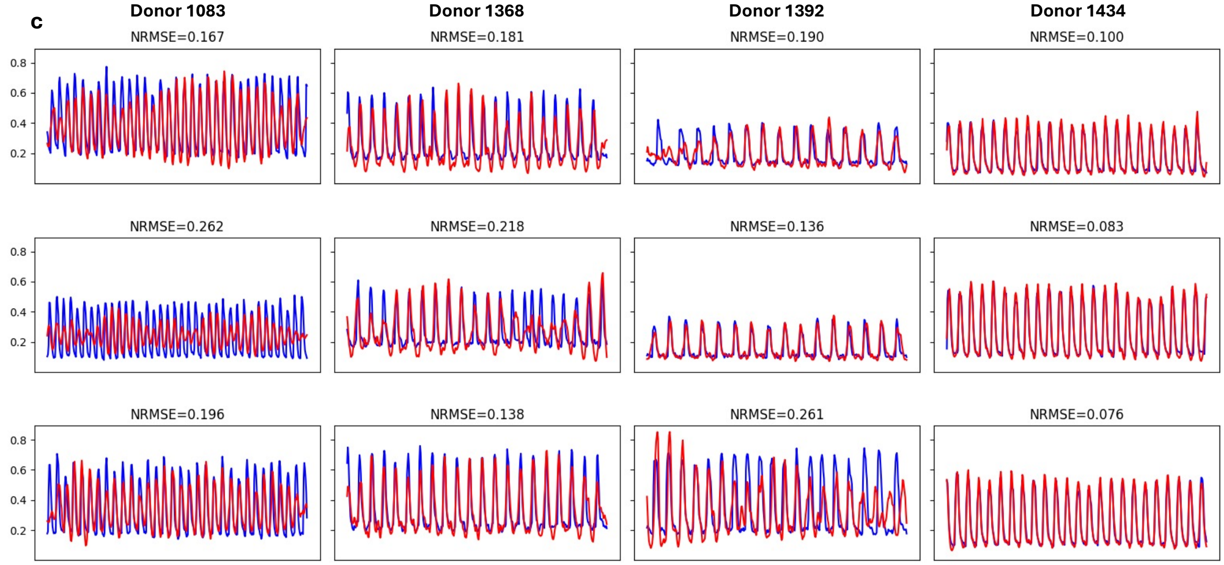


**Supplemental Figure 1. Autoencoder training performance and cross-validation.** (a) Training and validation loss curves over 500 epochs, indicating effective generalization without overfitting. (b) Five-fold cross-validation results showing highly consistent MSE values across all folds (train: 0.00547–0.00548; test: 0.00545–0.00550), confirming stable reconstruction performance across held-out data partitions. (c) Representative reconstructions of baseline calcium transient signals from the validation set, showing close agreement between original (blue) and reconstructed (red) traces.

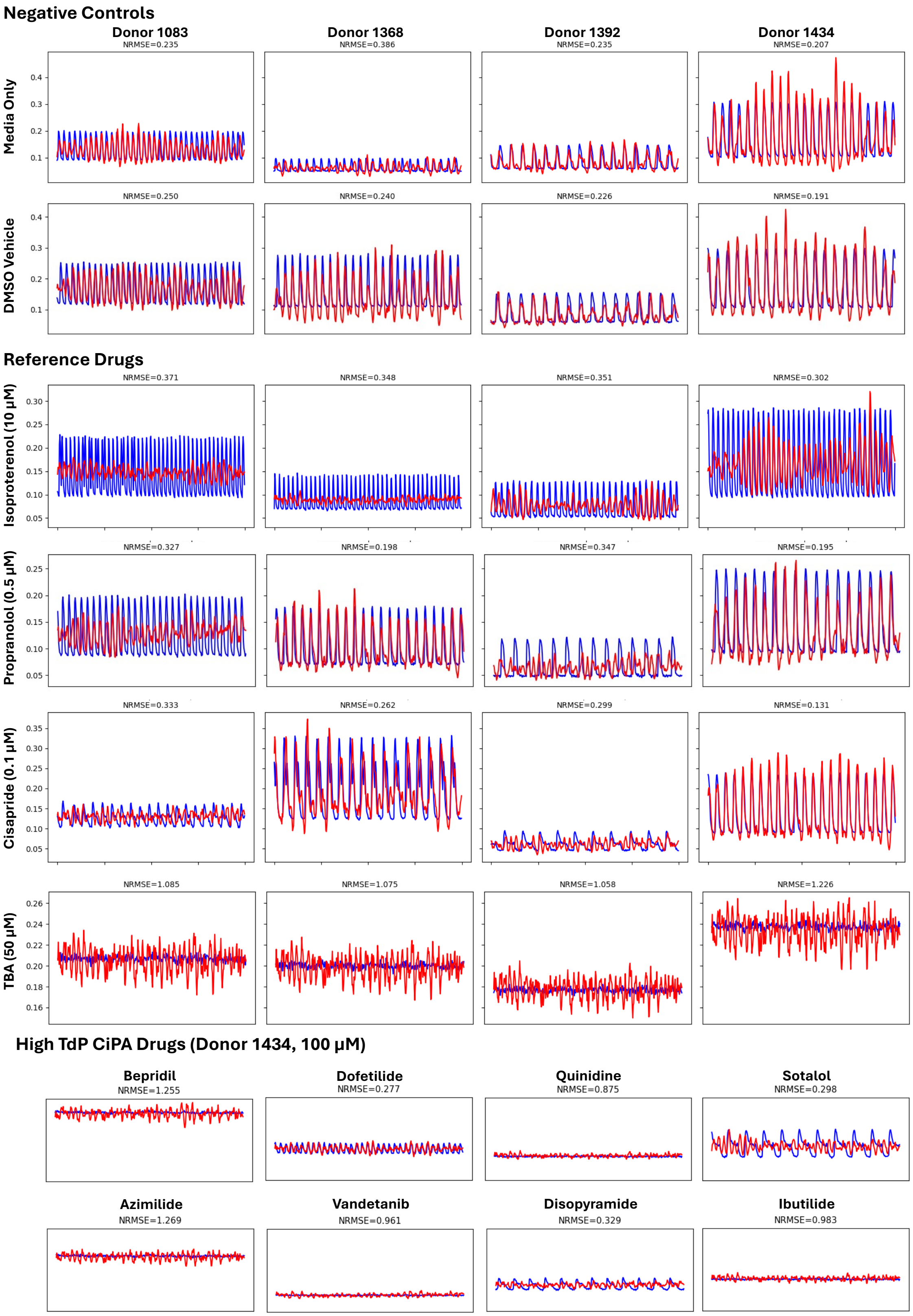


**Supplemental Figure 2. Reconstruction error profiles across reference groups.** Representative signal reconstruction for negative controls (media only and DMSO vehicle), reference drugs (Isoproterenol, Propranolol, Cisapride and TAB) and high-TdP CiPA drugs at highest concentration (100 µM). The reconstruction errors from these drugs were used to define cardiotoxicity thresholds.

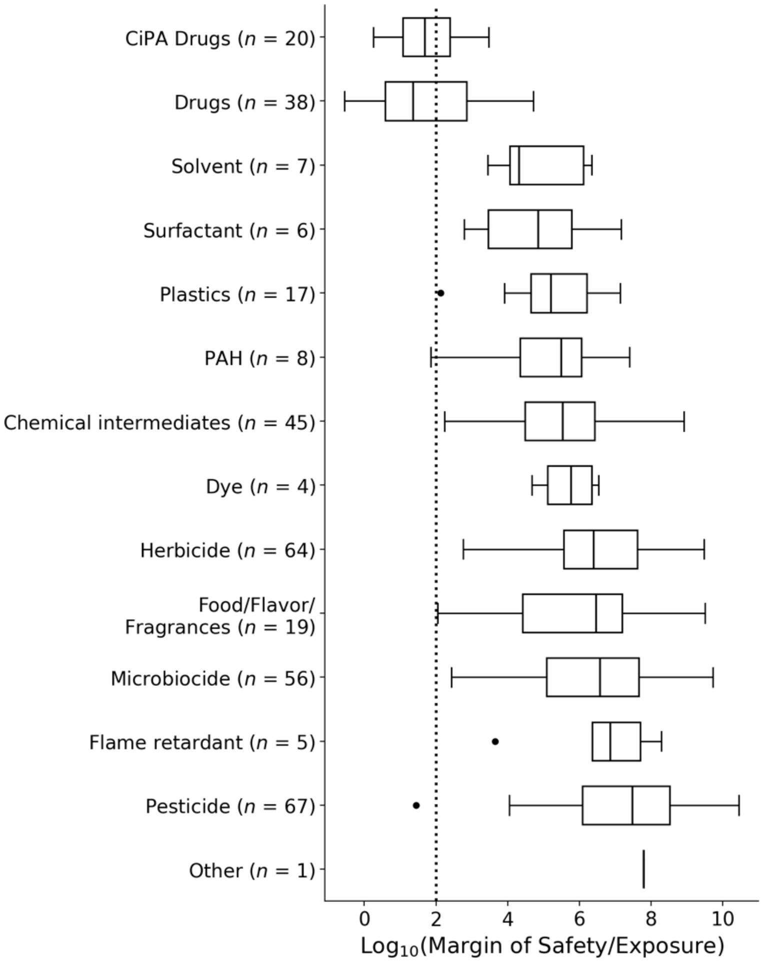


**Supplemental Figure 3.** Safety margins across chemicals with available exposure data, where pharmaceuticals were evaluated using population median $C_{max}$ values and environmental chemicals were evaluated using the median steady-state plasma concentrations derived from ExpoCast exposure estimates and toxicokinetic modeling

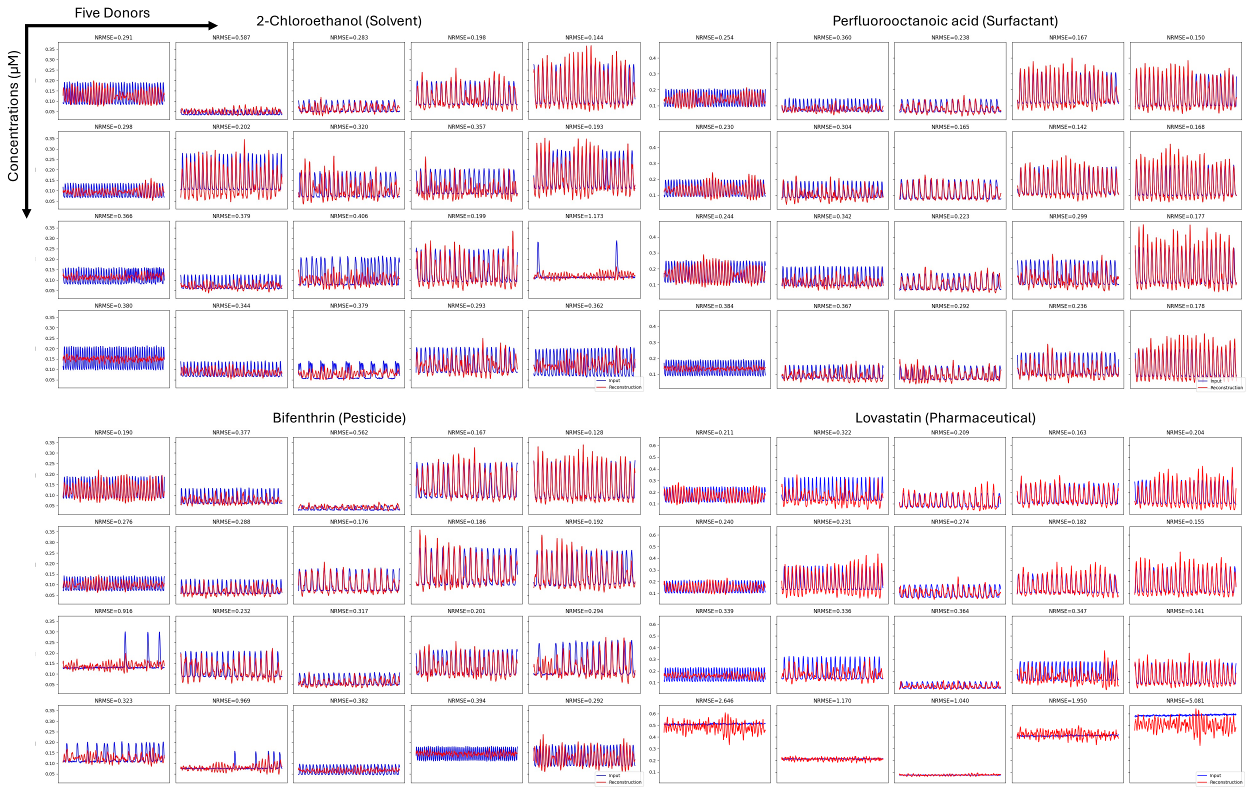


**Supplemental Figure 4.** Donor-specific drug response differences on representative ToxCast chemicals. Signal reconstruction from five donors, including held-out donor 1118, for four ToxCast chemicals at four concentrations. Inter-donor variability in NRMSE patterns reflects differences in cardiomyocyte sensitivity and calcium handling capacity across genetically distinct hiPSC-CM lines.

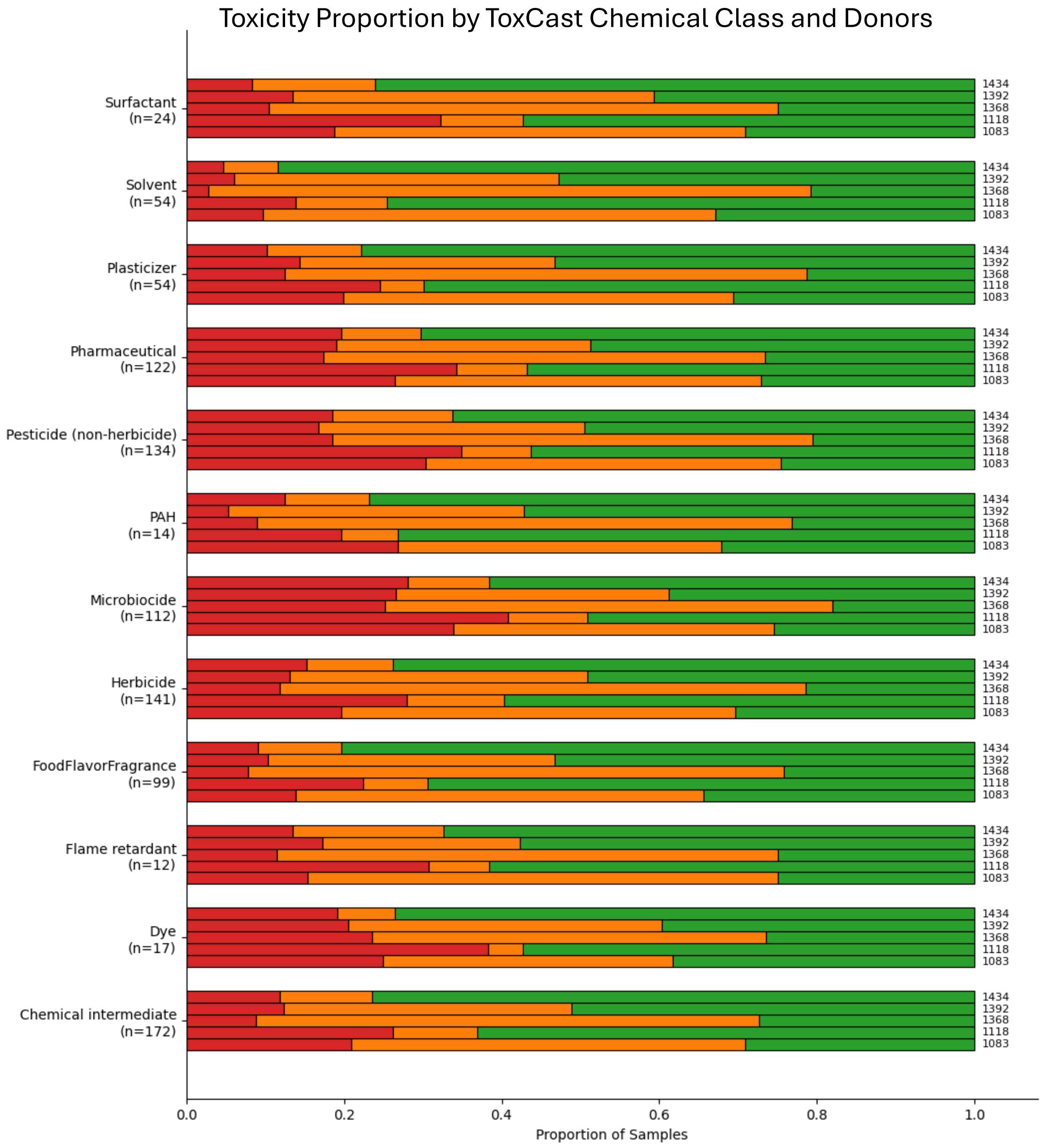


**Supplemental Figure 5. Donor-stratified toxicity proportion across ToxCast chemical classes.** The proportion of signals classified as low (green), medium (orange), and high (red) toxicity for each of the five donors (1434, 1392, 1368, 1118, 1083) within each ToxCast chemical class. Donor-specific patterns are visible across all chemical classes, with donor 1083 consistently showing elevated high-toxicity fractions and donor 1434 showing low-toxicity responses.

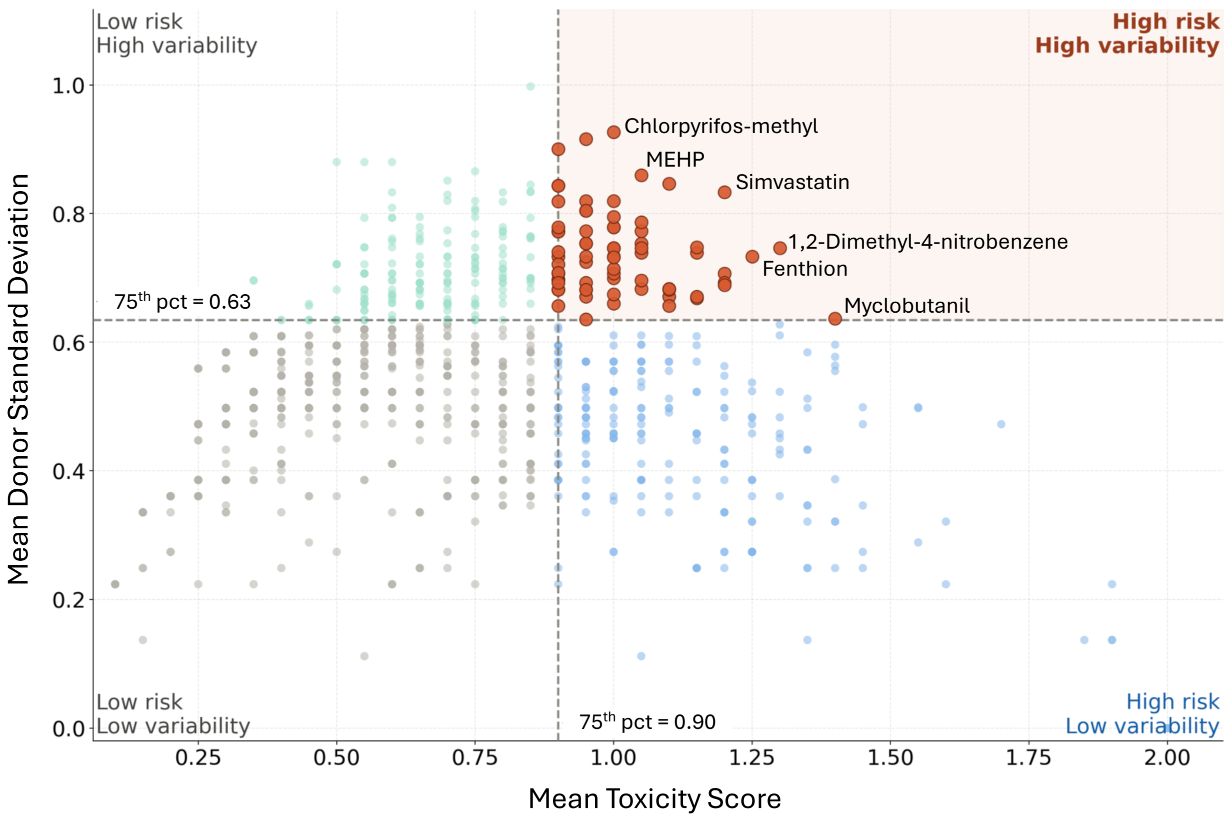


**Supplemental Figure 6.** Compound-level mean toxicity score versus mean donor standard deviation across the ToxCast library. Dashed reference lines indicate the 75th percentile thresholds for mean toxicity score (0.90) and mean donor STD (0.63), defining four quadrants. Representative chemicals in high risk/high variability quadrant (upper right, shaded) include Chlorpyrifos-methyl (pesticide, banned in European Union), MEHP (plasticizer from food packaging), Simvastatin (prescription drug for high cholesterol), 1,2-Dimethyl-4-nitrobenzene (chemical intermediate in industrial chemical production), Fenthion (pesticide), and Myclobutanil (fungicide).

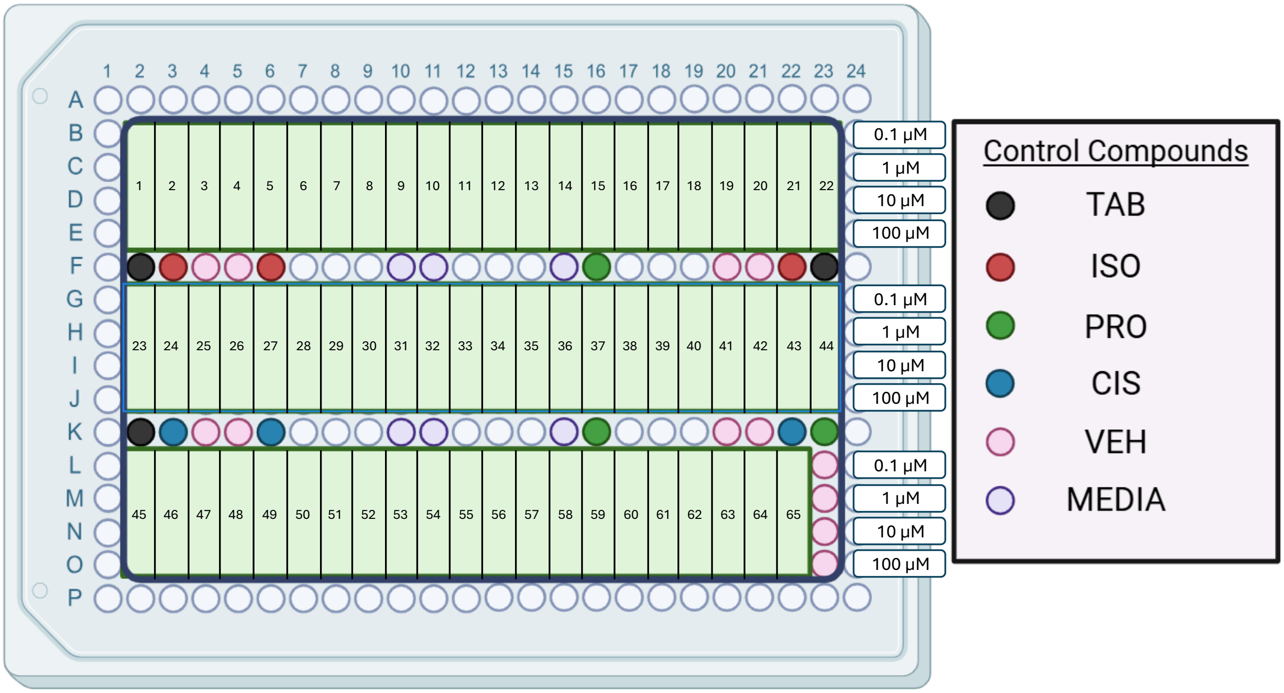


**Supplemental Figure 7.** Representative Plate Design. One plate in 384 format for high-throughput chemical screening. The plate contains 65 chemical compounds with 4 concentrations, as well as 6 control compounds.

**Supplemental Table 1.** Safety Margin and IC50 estimates for CiPA drugs using our autoencoder-based deep learning model

| COMPOUND | SAFETY MARGIN | IC50 (μM) | PLASMA CONCENTRATION [MM] |
|---|---|---|---|
| BEPRIDIL | 33.97 | 1.07 | 0.0315 |
| DOFETILIDE | NA | >100 | 0.0021 |
| QUINIDINE | 9.55 | 8.11 | 0.8492 |
| SOTALOL | 0.64 | 9.35 | 14.6864 |
| AZIMILIDE | 6.44 | 2.55 | 0.3960 |
| VANDETANIB | 2.65 | 11.27 | 4.2600 |
| DISOPYRAMIDE | 13.45 | 86.06 | 6.4000 |
| IBUTILIDE | 55.5 | 7.77 | 0.1400 |
| TERFENADINE | NA | NA | 0.0003 |
| ONDANSETRON | 32.63 | 11.77 | 0.3585 |
| ASTEMIZOLE | 144.87 | 1.13 | 0.0078 |
| CLARITHROMYCIN | 14.5 | 17.41 | 1.2000 |
| CLOZAPINE | 51.05 | 4.37 | 0.0856 |
| CISAPRIDE | 9,307.69 | 24.2 | 0.0026 |
| METOPROLOL | 96.625 | 38.65 | 0.4000 |
| CHLORPROMAZINE | 117.97 | 4.07 | 0.0345 |
| PIMOZIDE | 1,180.77 | 3.07 | 0.0026 |
| DOMPERIDONE | 224.4 | 11.22 | 0.0500 |
| DROPERIDOL | 8.49 | 3.16 | 0.3720 |
| RISPERIDONE | 10.15 | 0.41 | 0.0404 |
| MEXILETINE | 4 | 10 | 2.5032 |
| RANOLAZINE | 43.4 | 84.55 | 1.9482 |

**Supplemental Table 2.** Comparison of IC50 estimates for overlapped CiPA drugs between our calcium transient-based screening and patch-clamp studies on $Ca_{V1.2}$ ion channel.

| COMPOUND | AUTOENCODER | $CA_{V1.2}$[22] | $CA_{V1.2}$[23] |
|---|---|---|---|
| BEPRIDIL | 1.07 | 2.8 | 1.0 |
| DOFETILIDE | NA | NA | 26.7 |
| QUINIDINE | 8.11 | NA | 6.4 |
| SOTALOL | 9.34 | NA | 193.3 |
| TERFENADINE | NA | 0.7 | 0.9 |
| VERAPAMIL | 0.09 | 0.1 | 0.2 |

**Supplementary Table 3 (Separate Excel File).** IC50 estimated for all 1029 chemicals. 840 out of 1029 have defined IC50 values, and the rest with NA.

**Supplementary Table 4 (Separate Excel File).** Per-compound cardiotoxicity summary across 970 environmental chemicals with summary statistics by chemical class. Aggregate scores = sum of 5 donor votes (low = 0, medium = 1, high = 2) per dose (0.1, 1, 10, 100 μM). Risk tiers: High = max score ≥ 8; Medium = 5–7; Low < 5. Donor 1118 = held-out donor.
